# Rapid leaf trait response to growing-season meteorology in *Vitis:* Implications for leaf physiognomic paleoclimate reconstructions

**DOI:** 10.1101/706770

**Authors:** Aly Baumgartner, Michaela Donahoo, Daniel H. Chitwood, Daniel J. Peppe

## Abstract

**PREMISE OF THE STUDY:** The size and shape (physiognomy) of woody, dicotyledonous angiosperm leaves are correlated with climate and these relationships have been used to develop. proxies. These proxies assume that leaf morphology plastically responds to meteorological conditions and that leaf traits change isometrically through development.

**METHODS:** We used Digital Leaf Physiognomy (DiLP) to measure leaf characters of multiple *Vitis* species from the USDA Germplasm Repository in Geneva, NY from the 2012-2013 and 2014-2015 growing seasons. These growing seasons had different temperature and precipitation.

**KEY RESULTS:** We found three primary results: (1) there were predictable significant differences in leaf characters in leaves of different developmental stages along the vine, (2) there were significant differences in leaf characters in leaves of the same developmental stage between the growing seasons, and (3) there were significant differences in leaf characters between growing seasons.

**CONCLUSIONS:** We found that *Vitis* leaf shape had the strongest relationship with growing season meteorological conditions in taxa growing in their native range. In addition, leaves have variable phenotypic plasticity along the vine. We interpret that the meteorological signal was strongest in those leaves that have completed allometric expansion. This is significant for leaf physiognomic-paleoclimate proxies because these leaves are most likely to be preserved in leaf litter and reflect the type of leaves included in paleoclimate reconstructions. We found that leaf development does have the potential to be a confounding factor, but it is unlikely to exert a significant influence on analysis due to differential preservation potential.

## INTRODUCTION

The relationship between leaf size and shape (physiognomy) and climate in woody dicotyledonous angiosperms has been noted for over 100 years (Bailey and Sinnott, 1915, 1916) and has been used to develop proxies for reconstructing paleoclimate (e.g. Bailey and Sinnott, 1915, 1916; Wing and Greenwood, 1993; Wolfe, 1993; Wilf, 1997; Wilf et al., 1998; Jacobs, 1999, 2002; Gregory-Wodzicki, 2000; Adams et al., 2008; Peppe et al., 2011). Angiosperm leaves exhibit a dramatic diversity of sizes and shapes. Leaf shape is dynamic and changes across many scales, from the evolutionary timescales that differentiate species (Bailey and Sinnott, 1915, 1916; Schmerler et al., 2012), to phenotypic plasticity during the lifetime of a single plant (Royer et al., 2008, 2009; Royer, 2012b; Chitwood et al., 2015, 2016; McKee et al., 2019), to heteroblasty as a plant grows (Gould, 1993), to the allometric changes in a single leaf as it develops (Nicotra et al., 2011). Leaf physiognomic paleoclimate proxies are based on two tacit assumptions: (1) that plants plastically respond to changes in climate and (2) that changes in leaf physiognomy scale through growth and development. For decades, researchers have stated that a better understanding of phenotypic variation in leaves is necessary in order to understand its functional significance (e.g., Evans, 1972; Coleman et al., 1994). However, these assumptions have not been fully tested.

Changes on evolutionary timescales often lead to predictable patterns of leaf physiognomy; for example, high proportions of toothed leaves are found in cold temperate regions (Bailey and Sinnott, 1915, 1916), leaves tend to be smaller in arid environments and larger in wetter environments (e.g., Webb, 1968; Wilf et al., 1998; Jacobs, 1999; Malhado et al., 2009; Wright et al., 2017) and leaves tend to be more round in cooler environments and more elliptical in warmer environments (Peppe et al., 2011; Schmerler et al., 2012). Work by Royer et al. (2009) demonstrated that *Acer rubrum* exhibited rapid phenotypic plasticity during the lifetime of the plant when seeds were grown in a contrasting climate from their source. However, not all species exhibit phenotypic plasticity. For example, a study by Royer et al. (2008) demonstrated that while *Acer rubrum* exhibits phenotypic plasticity in response to changing temperature, *Quercus kelloggii* was phenotypically invariant to changing temperature. Further, species of the same genus may exhibit differences in temperature sensitivity (Royer et al., 2008; McKee et al., 2019) and the effects of temperature on leaf physiognomy are often species specific (McKee et al., 2019).

In addition to phenotypic plasticity, studies of leaf physiognomy must consider both allometric (differences in shape due to varying growth rates across an organ) and heteroblastic (differences in shape at successive nodes) influences on leaf shape (Chitwood et al., 2015, 2016). Leaf physiognomic paleoclimate proxies assume that leaf traits scale as a leaf matures (isometric change). However, this is often not the case. Phenotypic leaf traits change dramatically through growth and development (Evans, 1972). In the leaves of many dicotyledonous angiosperms, the maturation of xylem and phloem in the midrib and higher order venation proceeds from the lamina base to the tip (acropetal), while the maturation of smaller veins proceeds from the lamina tip to the base (basipetal) (Turgeon, 1989). The different directions of maturation lead to variation in expansion rates in different regions of the leaf (allometric change). Further, these patterns of allometric expansion are variable between species during leaf development (Das Gupta and Nath, 2015).

Heteroblasty is the difference in the shape of leaves from successive nodes due to shoot apical meristem development (Ashby, 1948; Jones, 1993, 1999; Poethig, 2010; Zotz et al., 2011). Heteroblastic differences can range from subtle to striking and are affected by both developmental and environmental factors (Poethig, 1990; Gould, 1993; Day et al., 1997; Kerstetter and Poethig, 1998; Winn, 1999; Darrow et al., 2001; Chitwood et al., 2012).

A drastic example is *Pseudopanax crassifolius* of New Zealand, which exhibits dramatically different leaves throughout its growth (Gould, 1193). Plasticity and heteroblasty can be easily confused and the terms are often used interchangeably (Diggle, 2002). Phenotypic plasticity refers to differences in leaf shape due to changing environmental conditions during the lifetime of a plant. However, it can be difficult to differentiate changes in leaf shape due to environmental phenotypic plasticity, allometric developmental changes or heteroblasty (Coleman et al., 1994).

*Vitis* is an economically important genus of temperate woody vines. The leaves of *Vitis* can range from simple to nearly compound, with shapes varying from orbicular to reniform to cordate (Chitwood et al., 2014). Previous work by Chitwood et al. (2016) analyzed >5,500 leaves from multiple species of *Vitis* and *Ampelocissus*. Their study showed that a more pronounced distal sinus, independent of species or developmental stage, was associated with a colder, drier growing season. In order to test the assumptions embedded in leaf physiognomic-paleoclimate proxies that leaf shape responds to meteorological changes and that leaf traits scale through development, we address 3 questions: Does *Vitis* leaf shape change isometrically or allometrically through development? Does *Vitis* leaf shape respond to changing temperature and precipitation between growing seasons? Are leaf shapes associated with developmental stages distinguishable from those due to meteorological influences? To address these questions, we tested the relationships between leaf physiognomic-development and leaf physiognomic-meteorological relationships in 4 species of *Vitis* collected in two growing seasons with different mean temperature and precipitation.

## MATERIALS AND METHODS

### Leaf collection

The leaves of four species of *Vitis* (*V. acerifolia, V. aestivalis, V. amurensis*, and *V. riparia*) were sampled from the USDA germplasm repository in Geneva, New York, USA (Fig. 1) (Chitwood et al., 2015, 2016). The leaves were collected from the same vines during June of the 2012-2013and 2014-2015 growing seasons and photographed for digital analysis. For a more detailed description of the collection process, see Chitwood et al. (2015) and Chitwood et al. (2016). The two growing seasons had different average temperature and precipitation: in 2012-2013 the mean annual temperature was 10.4 °C with a mean of 5.6 leaf wetness hours and in 2014-2015 the mean annual temperature was 8.2 °C with a mean of 3.8 leaf wetness hours. Following the methods of Chitwood et al. (2016), leaf wetness hours were used rather than mean annual precipitation because the pattern of precipitation was complicated (2014 had more precipitation than 2012, but 2013 had more precipitation than 2015), while leaf wetness hours showed a pattern of higher leaf wetness in 2012-2013than 2014-2015.

**Figure 1.**
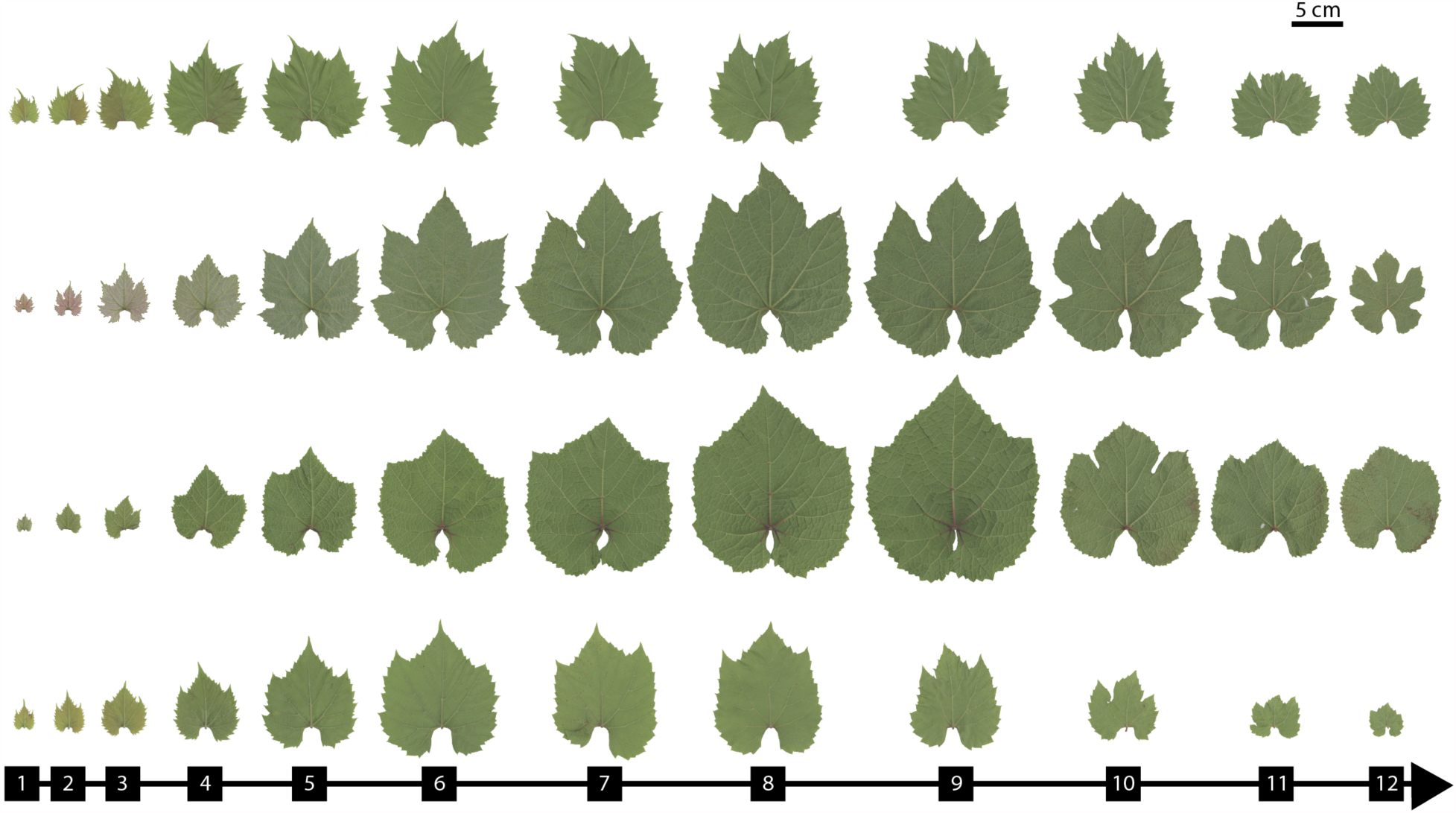
Four species of *Vitis*. From the top: *Vitis acerifolia, Vitis aestivalis, Vitis riparia*, and *Vitis amurensis*. Leaves are numbered from shoot tip (1) to base (12). Petioles are aligned to show differences in leaf size and shape between species and through development. The scale bar is 5 cm.

Phylogenetic analysis of *Vitis* places a *V. acerifolia, V. aestivalis*, and *V. riparia* in distinct, but closely associated clades, while *V. amurensis* is a phylogenetically distinct group (Zecca et al., 2012). *V. acerifolia, V. aestivalis*, and *V. riparia* are all native to North America. *V. riparia* has the widest range and spans from southern Canada to the Gulf of Mexico and from the Rocky Mountains to the Atlantic Ocean; the collection area is within its native geographic range. *V. aestivalis* spans from the northern United States to southern Texas and from Great Plains to the Atlantic Ocean, with the collection site being within its native geographic range. *V. acerifolia* spans from central Great Plains to northern Mexico and from the western Great Plains to the Gulf of Mexico. The collection area for this species is outside of its native geographic range. The native ranges of these species overlap in northern Texas and southern Oklahoma. *V. amurensis* is native to East Asia and spans from the Amur Valley of Russia to the Korean Peninsula and from eastern China to Japan. Thus, the collection site is far outside the geographic range of this species. The mean annual temperature and mean annual precipitation of the collection site are similar to those of the native range of *V. amurensis*; however, the mean annual range of temperatures is notably smaller (Qian and Ricklefs, 2004).

### Digital Leaf Physiognomy

Leaves were measured using the Digital Leaf Physiognomy (DiLP) protocol (Huff et al., 2003; Royer et al., 2005; Peppe et al., 2011) (Fig. 2). Images were prepared in Adobe Photoshop 8.0 (Adobe Systems Inc., San Jose, California, USA). Damaged leaf margins were reconstructed with a straight line; teeth were not reconstructed. When possible, the petiole was removed with a straight line cut at the intersection between the leaf lamina and the petiole. Primary teeth were defined as a vascularized extension of the margin supported by secondary or higher order venation; lobes were defined as vascularized extensions of the margin supported by primary venation (Wilf, 1997; Huff et al., 2003; Royer et al., 2005; Ellis et al., 2009). The leaf teeth were removed with a straight line cut from sinus to sinus. Secondary and tertiary teeth were treated as extensions of the superjacent primary tooth (Fig. 2).

**Figure 2.**
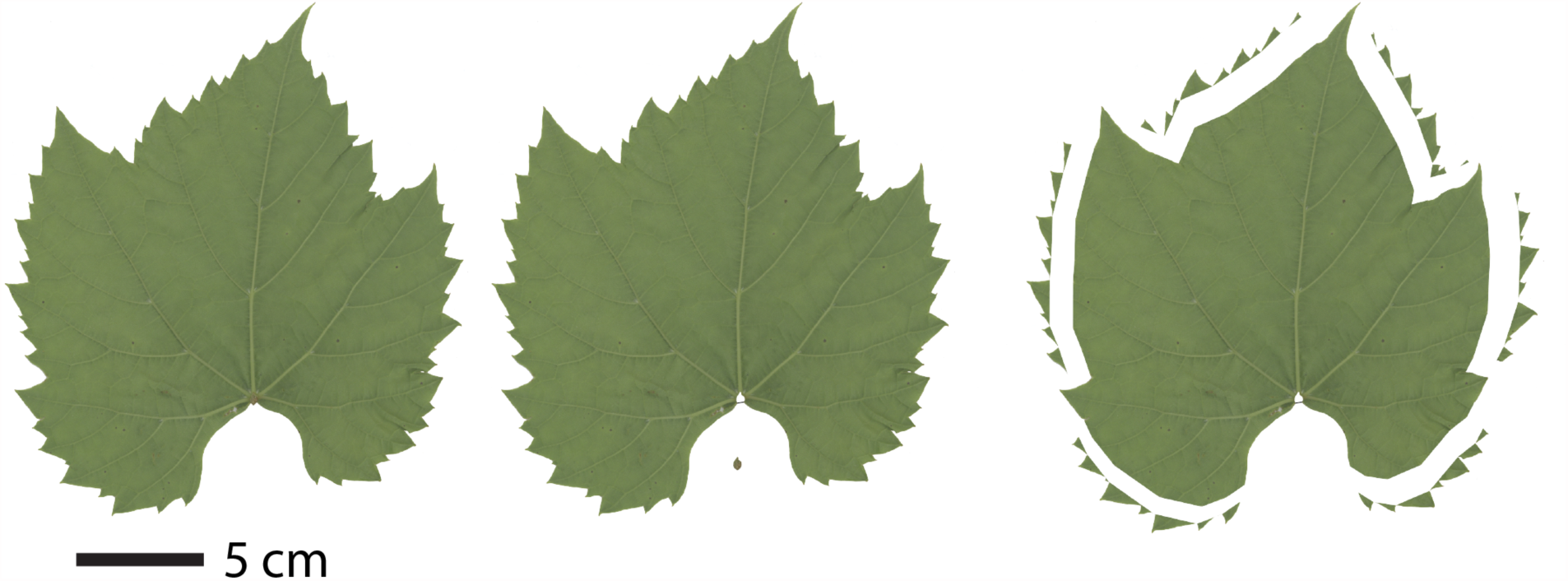
A *Vitis acerifolia* leaf processed with the Digital Leaf Physiognomy method. The petiole is removed in order to measure blade area. Internal perimeter is the perimeter of the leaf with the teeth removed.

Digital measurements were made using ImageJ (Abràmoff et al., 2004). Images were converted to greyscale and the threshold was adjusted so the leaf was a solid color and distinct from the background. Blade area, feret diameter, major feret, perimeter, internal perimeter, primary teeth, secondary teeth, total teeth, and tooth area were measured. See Royer et al. (2005) for a complete description of leaf characters used in DiLP.

### Morphometric Analysis, Statistics, and Visualization

To correct for differences in leaf size, ratios of leaf characters were used to compare between different species and leaf numbers. These ratios were feret diameter ratio, tooth area: perimeter, tooth area: internal perimeter, tooth area: blade area, total teeth: perimeter, total teeth: internal perimeter, total teeth: blade area, perimeter: area, perimeter ratio, compactness, and shape factor; total teeth, leaf area and average tooth area were also used to compare changes in leaf shape. Analyses of leaves were limited to the highest leaf number that had at least two leaves for both growing seasons and was at least twice as abundant in the dataset as the next highest leaf. *V. acerifolia* and *V. aestivalis* were limited to leaves 1-12, *V. amurensis* was limited to leaves 1-13, and *V. riparia* was limited to leaves 1-14 (Fig. 1). In this study, leaf number was measured from vine tip to vine base; leaf 1 is the youngest and leaf age increases with leaf number. For *V. acerifolia*, 11 plants were sampled with 101 leaves analyzed from 2013 and 102 leaves analyzed from 2015 (Table 1 and S1). For *V. aestivalis*, six plants were sampled with Ed leaves analyzed from 2013 and Ee leaves analyzed from 2015. For *V. amurensis*, 15 plants were sampled with 143 leaves analyzed from 2013 and 140 leaves analyzed from 2015. For *V. riparia*, 17 plants were sampled with 173 leaves analyzed from 2013 and 142 leaves analyzed from 2015.

**Table 1.** Plant and leaf totals from each species for 2013 and 2015.

| Species | Plants | 2013 | 2015 |
| --- | --- | --- | --- |
| *V. acerifolia* | 11 | 101 | 102 |
| *V. aestivalis* | 6 | 49 | 45 |
| *V. amurensis* | 15 | 143 | 140 |
| *V. riparia* | 17 | 173 | 142 |

Bootstrap forest analyses (JMP®, Version 12.0 SAS Institute Inc., Cary, NC) were performed to determine if the different *Vitis* species could be differentiated on the basis of the measured leaf characters, and if the leaves of each species could be differentiated by year. Student’s t tests were performed in R (base package) to determine whether leaf shape characters changed between growing seasons. Because developmental and environmental influences on leaf shape can be confounding, linear modeling (base package) and breakpoint analyses were performed (’segmented’, Muggeo, 2008) to determine whether there were consistent developmental “bins” of distinct leaf shapes. Leaf shape characters that were significantly different between the two growing seasons were analyzed separately. Once bins were determined, additional student’s t tests were run to determine whether changes in leaf shape were consistent through all bins. Normality was assumed. Finally, the changes between growing seasons were compared to the correlation table in Peppe et al. (2011) to determine whether changes in leaf shape were as expected for the known changes in temperature and precipitation between the growing seasons in 2012-2013 and 2014-2015.

## RESULTS

Bootstrap forest analysis showed that all four *Vitis* species could be distinguished based on leaf shape characters. The misclassification rate was 0.0521 (Table S2 and S3). Thus, all additional analyses were performed on each species individually. Bootstrap forest analysis also showed that for all species growing seasons were distinguishable based on the measured leaf characters (Table S4 and S5); the misclassification rate ranged from 0.0410-0.087 (Table S4).

All leaf shape characters changed along the vine (Fig. 3). Some characters were more sensitive to developmental and heteroblastic changes than others, and some characters demonstrated greater differences between growing seasons than others. For example, feret diameter ratio increased as leaves matured, meaning that young leaves were slightly more linear while mature leaves were nearly round. Characters related to the total number of teeth (total teeth: perimeter, total teeth: internal perimeter, and total teeth: blade area) were highly variable in the young, immature leaves but were conserved in mature leaves. Perimeter: area and compactness were also most variable in the young, immature but were conserved in mature leaves. Perimeter ratio and shape factor had inverse relationships with leaf number, and perimeter ratio decreased as leaves matured while shape factor increased as leaves matured. Tooth area: perimeter and tooth area: internal perimeter appeared to be driven by changes in tooth area, and tooth area: blade area was relatively stable in *V. acerifolia* and *V. riparia* but highly variable in *V. aestivalis* and *V. amurensis*. There were four patterns of leaf shape change (Fig. 3). First, through development, the average tooth area, feret diameter ratio, tooth area: perimeter, and tooth area: internal perimeter of the leaves of *V. acerifolia, V. aestivalis*, and *V. riparia* changed in a similar way, while *V. amurensis* displayed a slightly different developmental pattern (Fig. 3A). Second, for tooth area: blade area, perimeter ratio, compactness, and shape factor the leaves of some species had statistical differences between growing seasons and these different species clustered together based on growing season (Fig. 3B). Third, for total teeth: blade area, total teeth: perimeter, and total teeth: internal perimeter, the youngest leaves of all species were distinct, but older leaves were virtually indistinguishable (Fig. 3C). Finally, for total teeth and perimeter: area, there is no obvious pattern between growing seasons and all species had similar changes along the vine (Fig. 3D). Thus, leaf shape was plastic and changed between consecutive leaves of a single species from a single growing season. as well between leaves of a single species of the same leaf number between growing seasons.

**Figure 3.**
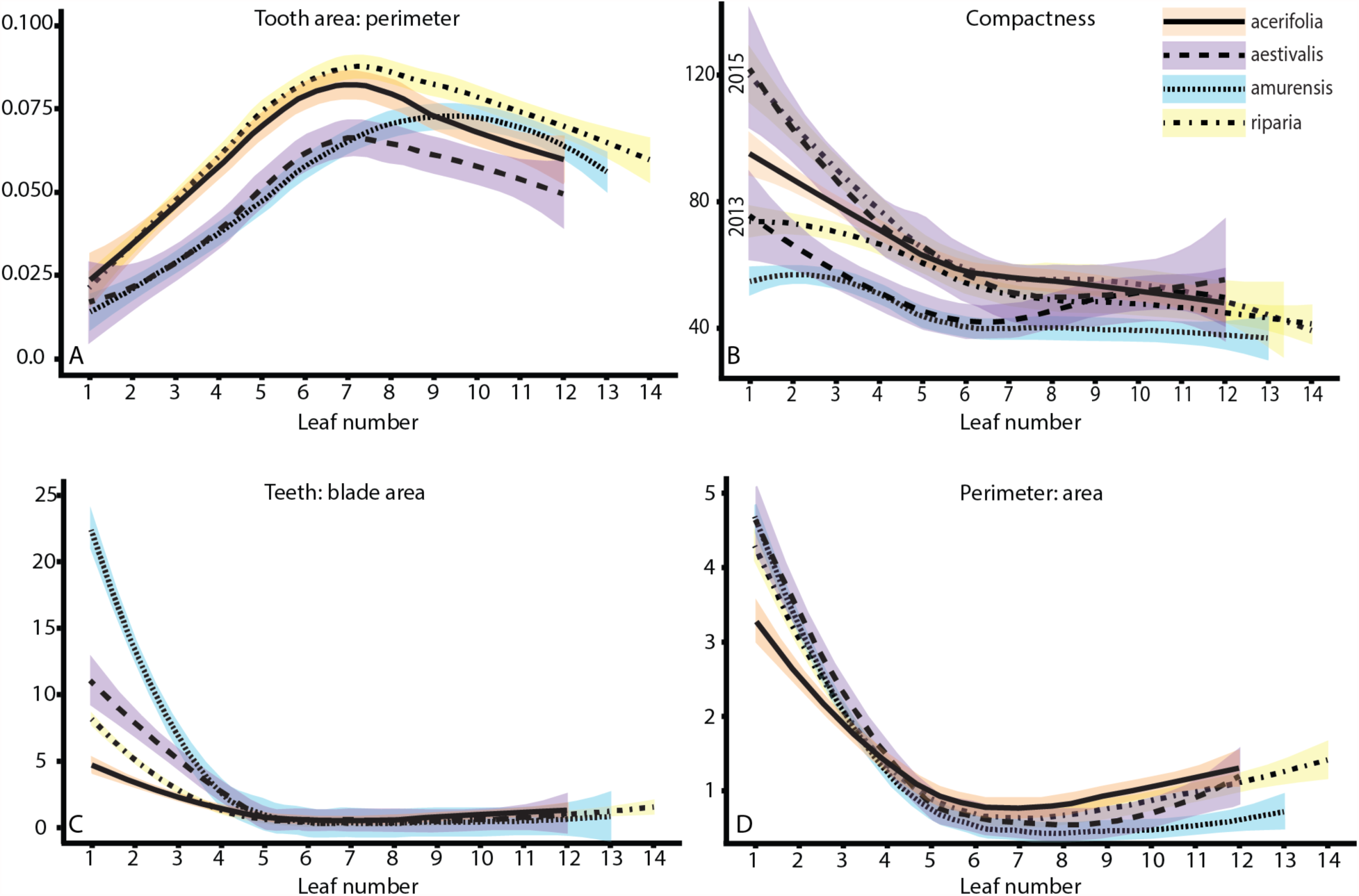
Patterns of leaf characters (A) were similar for North American species but not *V. amurensis*, (B) were different between growing seasons, (C) changed through development, and (D) were nonexistent. (A) The leaves of *V. acerifolia, V. aestivalis*, and *V. riparia* followed the same trend while *V. amurensis* values were offset, as demonstrated by tooth area: perimeter. (B) The leaves of some species had statistical differences between growing seasons and different species clustered together based on year, as demonstrated by compactness. (C) The youngest leaves of all species were distinct but older leaves were virtually indistinguishable, as demonstrated by total teeth: blade area. (D) Leaves showed no obvious pattern between growing seasons and all species showed similar changes through development.

Because species were distinguishable by growing season, we used student’s t tests to determine which aspects of leaf shape were plastic. All four species of *Vitis* had statistically significant differences between growing seasons for at least one leaf shape character: *V. acerifolia* showed differences in tooth area: blade area, *V. aestivalis* showed differences in total teeth, tooth area: blade area, and compactness, *V. amurensis* showed differences in total teeth, perimeter ratio, average tooth area, tooth area: perimeter, and tooth area: internal perimeter, and *V. riparia* showed differences in perimeter ratio, total teeth: perimeter, total teeth: internal perimeter, tooth area: blade area, tooth area: internal perimeter, compactness, and shape factor (Table S6, S7, S8, S9). All of the North American species *(V. acerifolia, V. aestivalis*, and *V. riparia)* also had statistical differences for tooth area: blade area.

Because there were notable changes in leaf shape characters along the vine through development, we used linear modeling to determine how much of the variance in leaf characters could be explained by leaf number (heteroblastic changes). The linear relationship between leaf shape and leaf number varied widely between the different species, though for all species more than 30% of the variance of compactness and shape factor could be explained by leaf number; in *V. aestivalis* this was only the case for the 2014-2015 growing season (Table S10, S12, S14, S16). If there were dramatic differences in the variance between growing seasons, it would often be much higher in the 2014-2015 but not the reverse. We then used breakpoint analysis to determine when changes in leaf shape characters occurred. This analysis showed three leaf forms: a distinct “young” form, a transitional “mid” form, and a distinct “old” form. The actual breakpoint differed for each variable and for each species; however, generally the “young” bin was leaves 1-2, the “mid” bin was leaves D-<c, and the “old” bin was leaves 11-12 (Table 2). We then used these shape bins to determine whether changes between growing seasons were driven by development or climate. The different species showed differential responses between growing seasons. In particular, *V. acerifolia* and *V. aestivalis* showed little plasticity in leaf shape between growing seasons. *V. acerifolia* only had significant differences in tooth area: blade area for the young and mid bins as well as the mean (Fig. 4), while *V. aestivalis* only had significant differences in total teeth for young and mid bins as well as the mean, for tooth area: blade area for young and mid bins as well as the mean, and for compactness and shape factor for the mid bin as well as the mean (Fig. 5). In contrast, *V. amurensis* and *V. riparia* showed considerable plasticity between growing seasons. *V. amurensis* had significant differences between growing seasons for tooth area: blade area, compactness and shape factor for the young bin, for leaf area, perimeter: area, and total teeth: blade area for young and mid bins, for total teeth and average tooth area for young and mid bins and the mean, for total teeth: perimeter and total teeth: internal perimeter for the mid bin, for tooth area: perimeter for the mid bin as well as the mean, for perimeter ratio for young and old bins as well as the mean, and for tooth area: internal perimeter for all bins as well as the mean (Fig. 6). *V. riparia* had significant differences between growing seasons for perimeter: area for the young bin, for tooth area: perimeter for young and mid bins, for total teeth: perimeter, total teeth: internal perimeter, compactness, and shape factor for young and mid bins as well as the mean, for perimeter ratio and tooth area: internal perimeter for the mid bin as well as the mean, and for tooth area: blade area for the mean (Fig. 7).

**Table 2.**
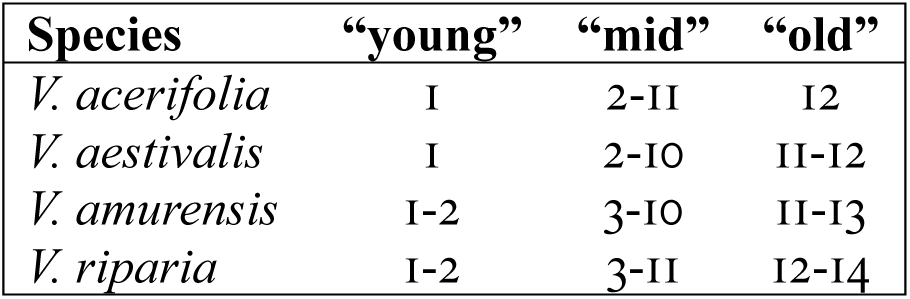
Breakpoint analysis of *Vitis acerifolia*, *Vitis aestivalis*, *Vitis amurensis*, and *Vitis riparia* based on all measured leaf shape characters.

**Figure 4.**
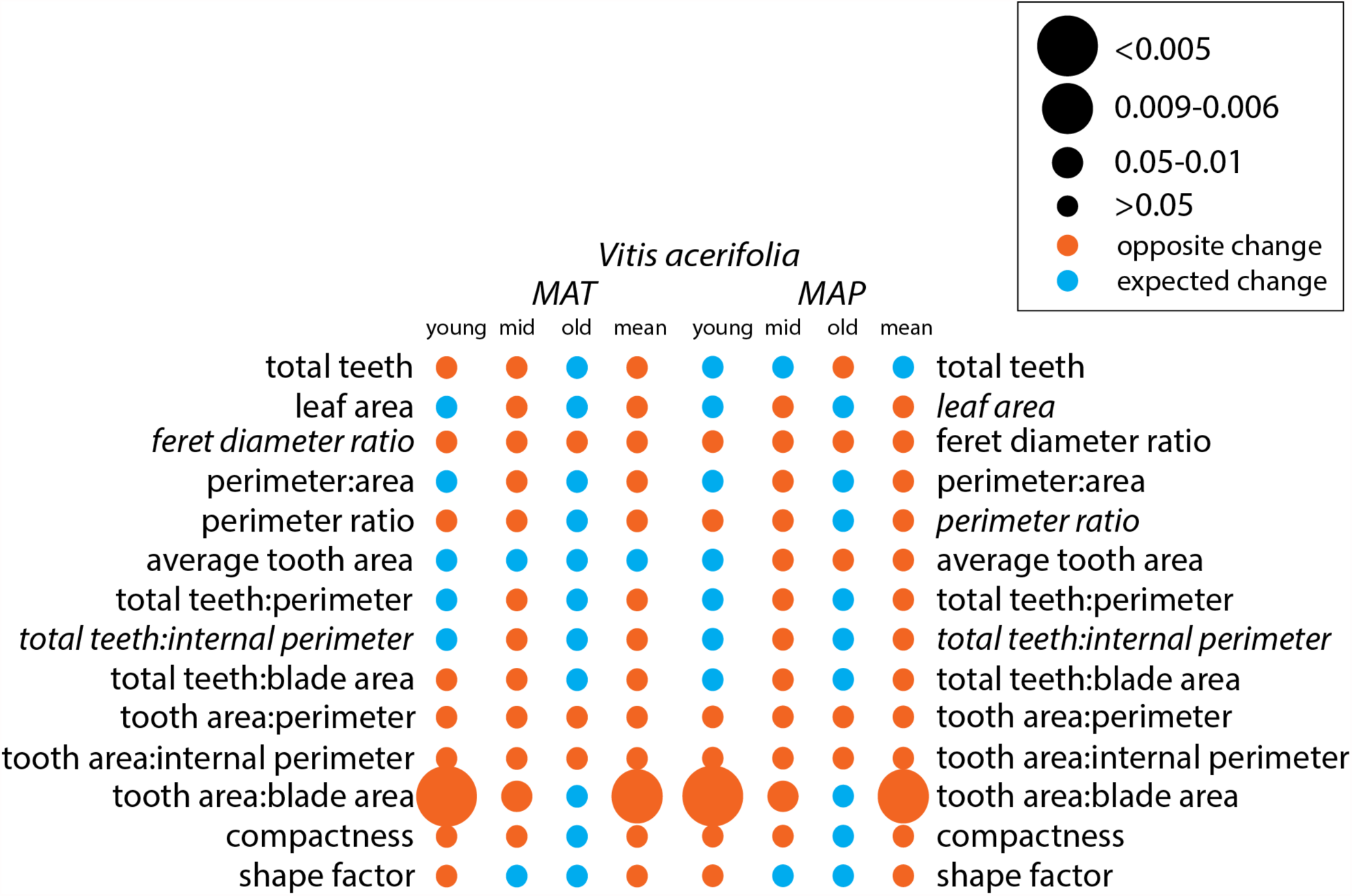
*Vitis acerifolia* binned significant effect table for mean annual temperature (MAT) and mean annual precipitation (MAP). Leaf shape variables included in the Digital Leaf Physiognomy climate equation are italicized. The larger the circle, the stronger the statistical relationship. Circle color is based on whether the change between growing seasons was as expected based on the correlation in Peppe et al. (2011) (blue) or the opposite of the expected change (orange).

**Figure 5.**
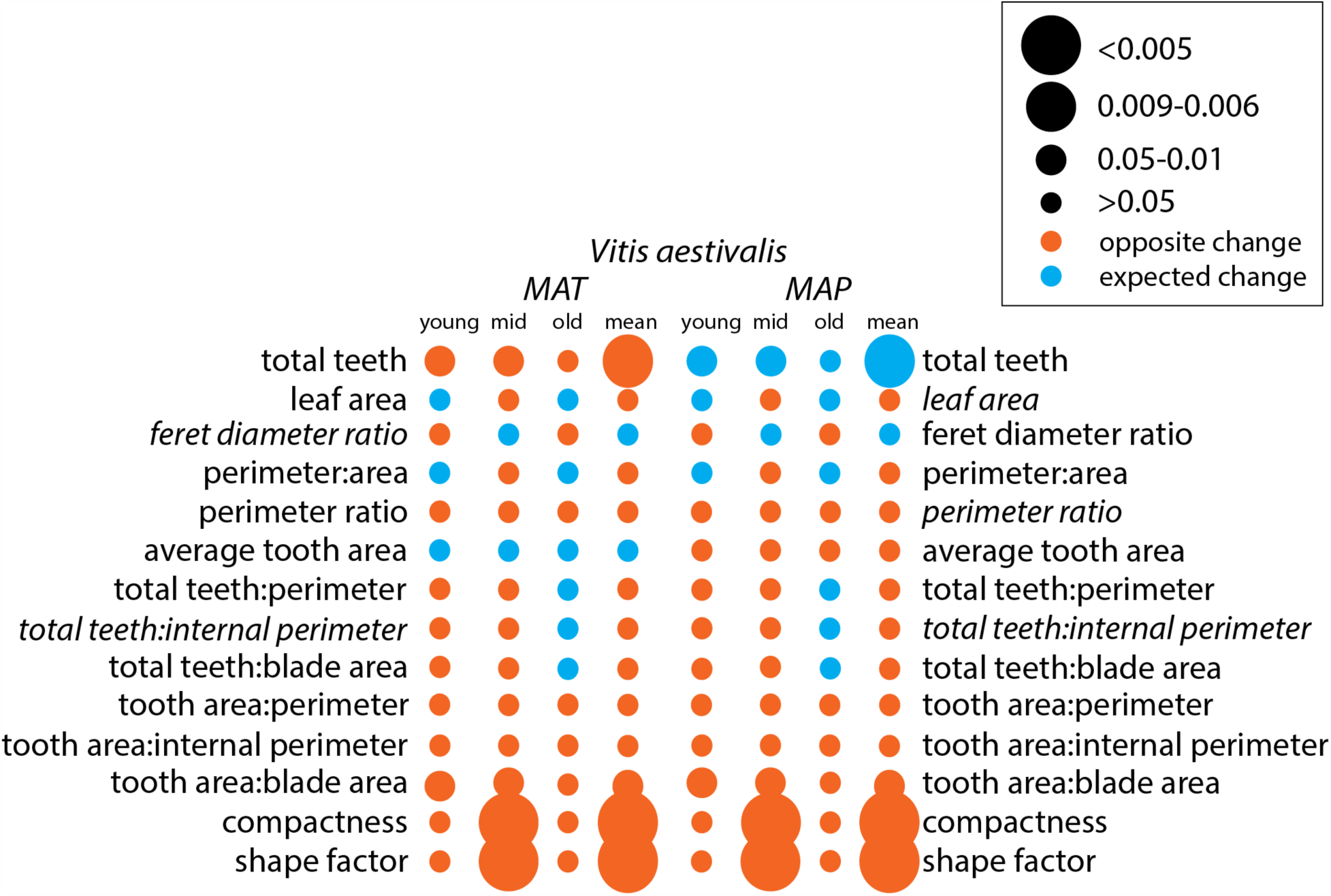
*Vitis aestivalis* binned significant effect table for mean annual temperature (MAT) and mean annual precipitation (MAP). Leaf shape variables included in the Digital Leaf Physiognomy climate equation are italicized. The larger the circle, the stronger the statistical relationship. Circle color is based on whether the change between growing seasons was as expected based on the correlation in Peppe et al. (2011) (blue) or the opposite of the expected change (orange).

**Figure 6.**
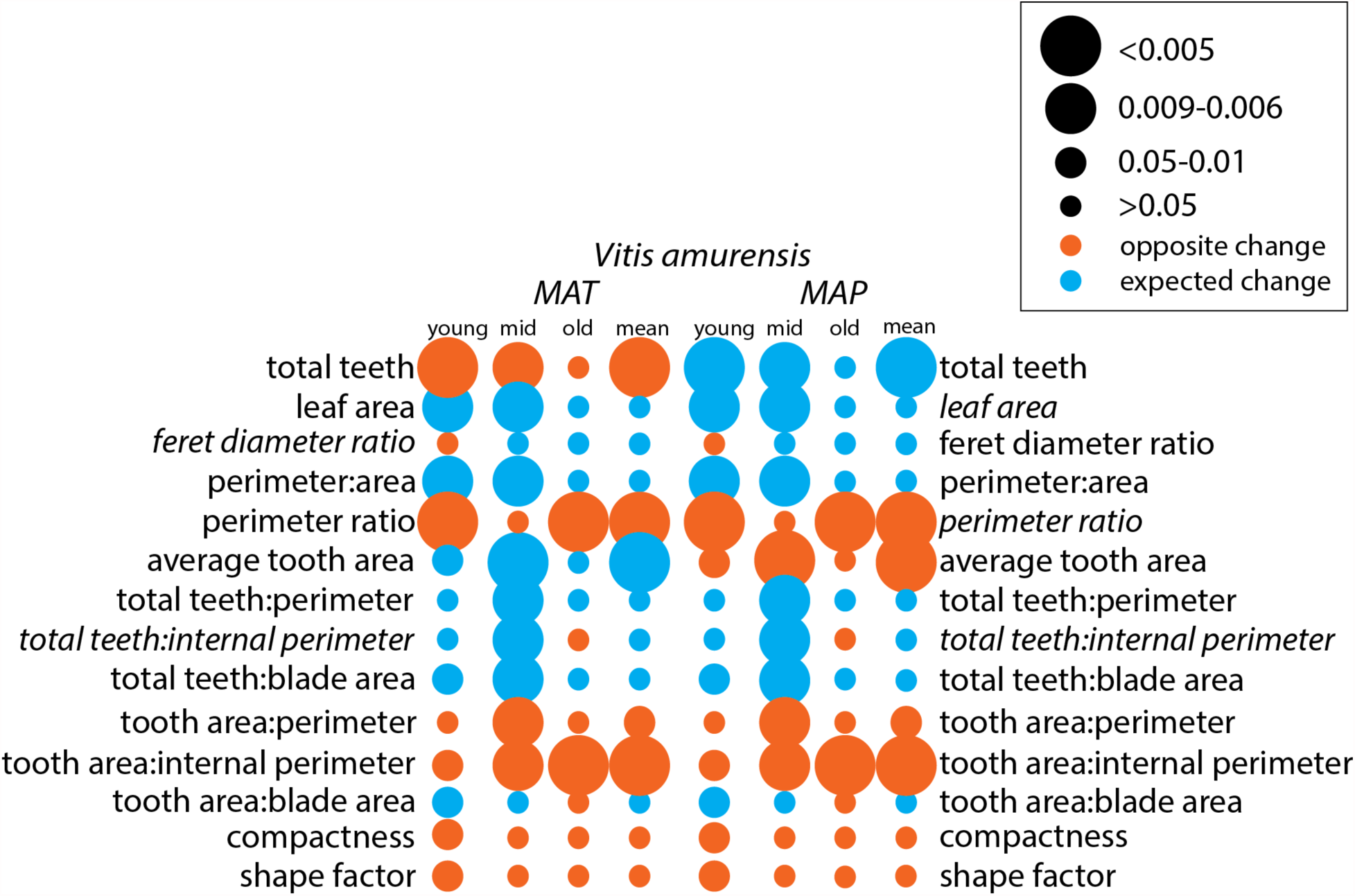
*Vitis amurensis* binned significant effect table for mean annual temperature (MAT) and mean annual precipitation (MAP). Leaf shape variables included in the Digital Leaf Physiognomy climate equation are italicized. The larger the circle, the stronger the statistical relationship. Circle color is based on whether the change between growing seasons was as expected based on the correlation in Peppe et al. (2011) (blue) or the opposite of the expected change (orange).

**Figure 7.**
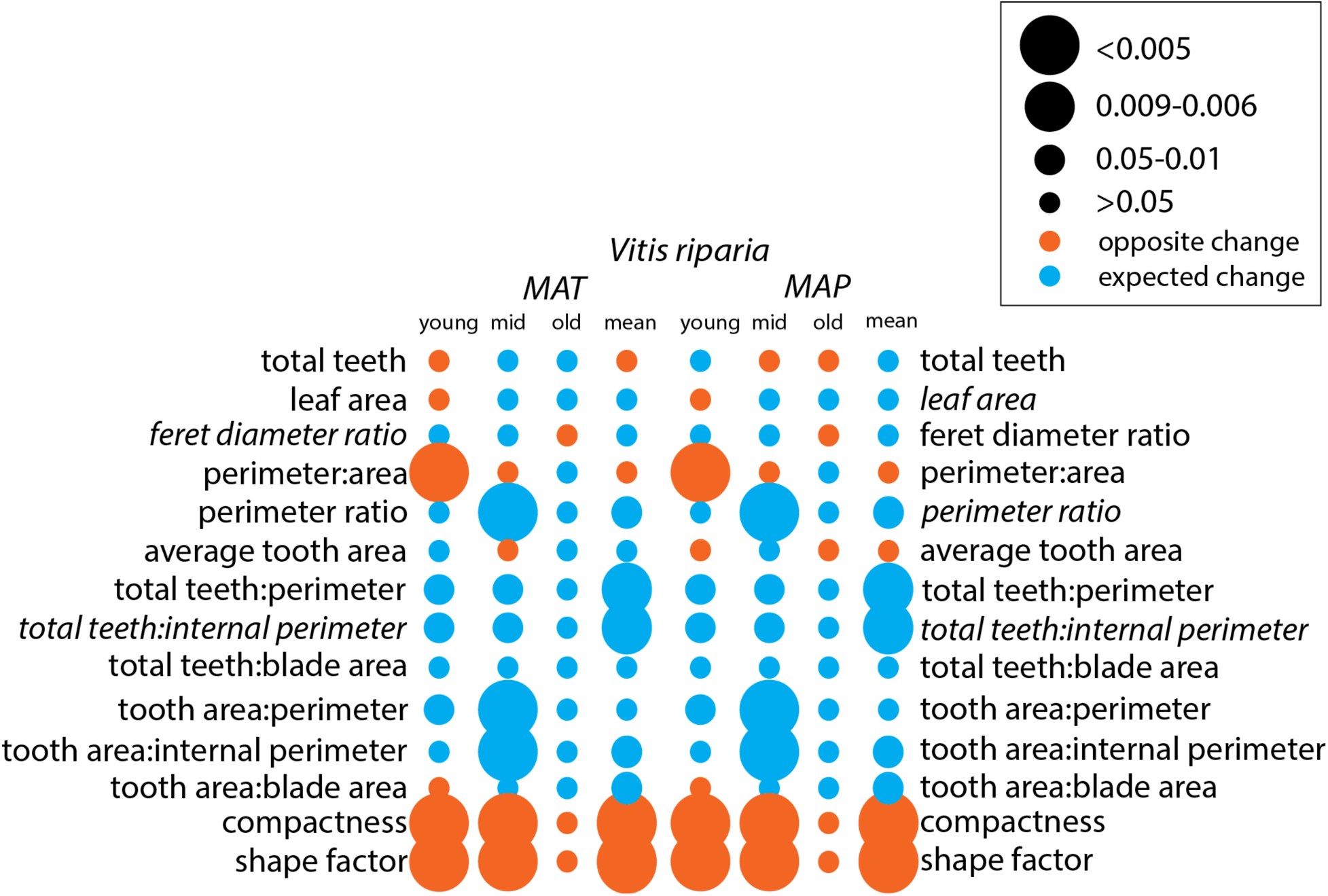
*Vitis riparia* binned significant effect table for mean annual temperature (MAT) and mean annual precipitation (MAP). Leaf shape variables included in the Digital Leaf Physiognomy climate equation are italicized. The larger the circle, the stronger the statistical relationship. Circle color is based on whether the change between growing seasons was as expected based on the correlation in Peppe et al. (2011) (blue) or the opposite of the expected change (orange).

After assessing for significant changes in leaf shape between growing seasons, we assessed how the leaf characters changed to determine if these differences were consistent with our expectations of phenotypic plasticity in response to changes in temperature and precipitation based on the DiLP model (Peppe et al., 2011). For all species, the correlation of the mean for each leaf shape character reflected the significant correlations derived from the analyses of each of the developmental bins. *V. acerifolia* and *V. aestivalis* were not sensitive to climate variability. For *V. acerifolia* only tooth area: blade area varied between growing seasons but did not change as expected with respect to temperature or precipitation (Fig. 4). For *V. aestivalis*, tooth area: blade area, compactness, and shape factor varied between growing seasons, but did not change as expected with respect to temperature or precipitation; total teeth changed as expected with respect to precipitation but not to temperature (Fig. 5). *V. amurensis* and *V. riparia* were sensitive to climate variability. For *V. amurensis*, perimeter ratio, tooth area: perimeter, and tooth area: internal perimeter varied between growing seasons, but did not change as expected with respect to temperature or precipitation; average tooth area changed as expected with respect to temperature but not to precipitation; and total teeth changed as expected with respect to precipitation but not to temperature (Fig. 6). For *V. riparia*, compactness and shape factor varied between growing seasons but did not change as expected with respect to temperature or precipitation, while perimeter ratio, total teeth: perimeter, total teeth: internal perimeter, tooth area: internal perimeter, and tooth area: blade area changed as expected with respect to both temperature and precipitation (Fig. 7).

## DISCUSSION

### Ontogenetic and Heteroblastic Physiognomic Change

Studies of leaf physiognomy must consider both allometric and heteroblastic influences on leaf shape (Chitwood et al., 2015, 2016). In addition to temperature and precipitation, the age of the vines changed between the growing seasons; however, *Vitis* is a long-lived woody perennial, so differences in leaf shape due to age of the plant were negligible compared to differences due to temperature and precipitation. The oldest leaves—at the base of the shoot—have reached maturity and are no longer expanding but, due to genetic constraints, are smaller than many of the younger leaves. The youngest leaves—at the tip of the shoot—are also relatively small but this is because they are still undergoing allometric expansion and have not yet reached maturity. In addition, the growth habit of *Vitis* allows us to see a snapshot of development, but also emphasizes heteroblastic differences. Differences in leaf shape give us a physiognomic roadmap of allometric changes (Fig. 1). Immature leaves have long narrow teeth, which gives the leaf an overall more linear shape. As leaves mature, the teeth become larger and more triangular. In addition, mature leaves have more pronounced basal lobes, which gives the leaf a more circular shape. These patterns in leaf shape reflect the underlying developmental signal.

Ontogenetic and heteroblastic changes in leaf shape were affected by changing meteorological conditions. For example, *V. acerifolia* only exhibited differences in leaf shape between growing seasons for tooth area: blade area. However, this difference was most pronounced in the youngest leaves and was less pronounced or entirely absent in the older, mature leaves (Fig. 4). *V. amurensis* had differences in tooth area: blade area, compactness and shape factor, but only in the youngest leaves (Fig. 6). It is worth noting that, with the exception of *V. amurensis*, the oldest leaves were invariant and not sensitive to changes in temperature or precipitation. This may mean that these leaves are optimized for early season growth and because *V. amurensis* was cultivated outside of its native geographic range, it is not optimized for North American spring. However, *V. acerifolia* was also cultivated outside of its native geographic range and did not exhibit phenotypic plasticity in the oldest leaves but this could be because *V. acerifolia* was not sensitive to changes in temperature.

### Climatic Physiognomic Change

In *Vitis*, leaf shape is patterned within buds during the year prior to budburst, therefore temperature and precipitation data were evaluated starting in the year before collection when temperatures began to rise above freezing until the time of collection (Carmona et al., 2008; Chitwood et al., 2016). Temperatures were consistently higher during the 2012-2013 growing season than 2014-2015 (2.2 °C). While the pattern of cumulative precipitation is more complicated, leaf wetness hours, which is influenced by precipitation in addition to other environmental factors including solar radiation, wind, and relative humidity, were higher during the 2012-2013 growing season than 2014-2015 (1.8 leaf wetness hours). Thus, the 2012-2013 growing was generally warmer and wetter than 2014-2015.

Previous work on *Vitis* by Chitwood et al. (2016) showed significant differences in the degree of dissection of the distal sinus between the 2012-2013 and 2014-2015 growing seasons; they were able to match the leaf to the growing season with 65.5-69.1% accuracy. The pronounced dissection of the distal sinus in the 2014-2015 growing season is associated with cooler and drier growing conditions. If physiognomic sensitivity in *Vitis* were driven by temperature, we would expect the leaves from the 2012-2013 growing season to have fewer, smaller teeth than the 2014-2015 growing season. If physiognomic sensitivity were driven by precipitation, we would expect the leaves from the 2012-2013growing season to be larger and less dissected with a greater number of larger teeth than the 2014-2015 growing season. Based on the correlations between the change in leaf shape and climate, these changes did not appear to be driven solely by temperature or precipitation, but a mixture of the two (Figs. 4–7). Overall, differences in leaf shape between growing seasons were primarily driven by tooth area and perimeter; to a lesser degree differences were driven by the total number of teeth and internal perimeter. Leaf area did not have a strong influence on differences in leaf physiognomy. When compared to the correlations from Peppe et al. (2011), differences in characters related to the total number of teeth were often correlated as expected with precipitation while differences related to tooth area were often correlated as expected with temperature.

*V. acerifolia, V. aestivalis, V. amurensis*, and *V. riparia* varied considerably in their phenotypic plasticity, however previous studies have shown that members of the same genus may have difference degrees of phenotypic plasticity and that effects of changing temperature on leaf physiognomy are species specific (Royer et al., 2008; McKee et al., 2019). It is also important to note that *V. acerifolia*, which had the fewest statistically significant differences in leaf traits between growing seasons, was the only species native to a warmer climate; *V. aestivalis* and *V. riparia* are both native to New York. Though *V. amurensis* experiences similar temperatures in its native range in east Asia, east Asian winters tend to be much colder than those in eastern North America while precipitation is generally higher in eastern North America than in east Asia (Qian and Ricklefs, 2004). All three North American taxa exhibited significant differences in tooth area: blade area. However, neither *V. acerifolia* nor *V. aestivalis* exhibited the expected change for either temperature or precipitation, while *V. riparia* exhibited the expected change for both.

*V. riparia* and *V. amurensis* were most sensitive to changes in temperature and precipitation, however each species had a different relationship between leaf shape and climate (Fig. 3). For example, both species had differences in perimeter: area in the youngest leaves, but *V. amurensis* exhibited the expected change for both temperature and precipitation while *V. riparia* did not exhibit the expected change for either. Conversely, both species had differences in tooth area: internal perimeter in the majority of their leaves, but *V. riparia* exhibited the expected change for both temperature and precipitation while *V. amurensis* did not exhibit the expected change for either. Compactness and shape factor were plastic in *V. aestivalis, V. amurensis*, and *V. riparia* but no species exhibited the expected changes for either temperature or precipitation. This suggests that either *Vitis* has a different relationship between compactness and shape factor and climate or that changes in compactness and shape factor are influenced by something other than temperature or precipitation in *Vitis*.

### Implications for Leaf Physiognomic Paleoclimate Reconstructions

Leaf physiognomic paleoclimate models are based on the assumption that leaves reflect their environment at the time of deposition. This relies on changes in leaf shape on both evolutionary and plant lifespan timescales. *Vitis* is a woody dicot angiosperm, and thus in theory can be used to estimate paleoclimate, but lianas have been shown to have a weaker relationship with leaf margin state and climate than trees or shrubs (Royer et al., 2012). In addition, *Vitis*’ preference for lowland and riparian environments is another potential confounding factor that can influence paleoclimate estimates (see discussion in Royer, 2012a).

Previous work on *Vitis* has shown that members of the genus are climatically sensitive (Chitwood et al., 2016). However, not all species of *Vitis* exhibited phenotypic plasticity in the variables of the DiLP paleoclimate equations (feret diameter ratio and total teeth: internal perimeter for MAT and leaf area, perimeter ratio, and total teeth: internal perimeter for MAP). *V. acerifolia* and *V. aestivalis* did not have significant differences in any of the DiLP variables and no species had any differences in feret diameter ratio. *V. amurensis* had significant differences in total teeth: internal perimeter and leaf area for some leaves but neither of these differences were significant at the species mean level; perimeter ratio was significantly different at the species mean level. *V. riparia* only had significant differences in total teeth: internal perimeter and perimeter ratio.

Interestingly, species means did not reflect the differences in phenotypic plasticity along the vine. For example, *V. riparia* exhibited small (though significant) differences in total teeth: perimeter and total teeth: internal perimeter in the young and mid bins, however the species mean showed large differences (Fig. 7). Similarly, there were no differences in tooth area: blade area in any bin but the species mean showed a small but significant difference. *V. amurensis* exhibited significant differences in total teeth: perimeter, total teeth: internal perimeter, and total teeth: blade area in the mid bin but none of these characters were significant at the species mean level (Fig. 6). This is significant because leaf physiognomic paleoclimate methods rely on species means to estimate paleotemperature and paleoprecipitation. These means use all data and are therefore a good summary, however the results here suggest that a grand species mean may not fully reflect the variability within a taxon and mask any variability that occurs through development.

These results suggest that *Vitis* leaf shape has the strongest relationship with climate in taxa growing in their native range. In addition, leaves have variable phenotypic plasticity along the vine, which could reflect the different roles of leaves through their ontogeny. The oldest leaves are invariant, which may be in order to maximize early season productivity. Physiognomic changes in newly flushed leaves are driven by allometric expansion, however it is unclear from this study whether the degree of climate sensitivity in newly flushed leaves varies through the growing season. In this analysis, the leaves of *V. amurensis* and *V. riparia*, the only species to exhibit climate sensitivity, that had completed most of their allometric expansion but did not flush early in the growing season (“mid” bin) had the most phenotypic plasticity. Therefore, we interpret that the climate signal was strongest in these leaves. This is significant for leaf physiognomic paleoclimate proxies because these leaves are most likely to be preserved in leaf litter and reflect the type of leaves included in paleoclimate reconstructions (Burnham et al., 1992). This suggests that while leaf development does have the potential to be a confounding factor, it is unlikely to exert a significant influence on analysis due to the unlikelihood of newly flushed or early season leaves being preserved.

## CONCLUSIONS

Leaf physiognomic paleoclimate proxies assume that the leaves of woody dicotyledonous angiosperms change isometrically through development and reliably reflect temperature and precipitation. We found that the leaves of four species of *Vitis* changed allometrically through development and that leaves had variable phenotypic plasticity along the vine. This suggests that, at least in species that demonstrate phenotypic plasticity, leaves reflect the meteorological conditions during bud patterning. In addition, the relationship between leaf shape and meteorological signal was strongest in leaves that had completed allometric expansion and in taxa growing in their native range. Finally, leaf development has the potential to be a confounding factor in leaf physiognomic paleoclimate proxies, but it is unlikely to exert a significant influence on analysis due to differential preservation potential. This is significant because these leaves are most likely to be preserved in leaf litter and reflect the most common type of leaves included in paleoclimate reconstructions. However, remaining questions include whether this pattern holds in species with different growth habits (e.g. shrubs, trees, etc.) or within a single species across its range.

## ACKNOWLEDGEMENTS

This work was supported by Baylor University Department of Geosciences’ Graduate Research Grant (AB). The authors thank J. Tubbs for his statistical assistance and J. White for his valuable suggestions. Additional thanks to A. Flynn, J. Milligan, J. Wagner, and G. Pruett for support and insight.

## AUTHOR CONTRIBUTIONS

A.B., D.C. and D.P. developed the project. D.C. collected the data, which was processed and analyzed by A.B. and M.D.. A.B., D.C. and D.P. developed the statistical analyses. A.B. wrote the manuscript with input from D.C., M.D. and D.P.

## DATA ACCESSIBILITY STATEMENT

The dataset analyzed during the current study are available in the Texas Data Repository (https://dataverse.tdl.org/dataverse/VitisLeaf).

